# Identification of *Nocardia* species using Autof MS 1000 and Bruker MALDI-TOF MS systems: A comparison

**DOI:** 10.1101/2021.01.11.426310

**Authors:** Bing Ma, Yunqi Tian, Yungang Han, Lifang Ban, Junwen Yang, Xinkun Qi, Changcheng Guo, Baoya Wang, Wenjuan Yan, Qiong Ma, Shanmei Wang, Youhua Yuan, Yi Li

## Abstract

*Nocardia* is an important cause of clinically invasive disease, but for most clinical laboratories, identification of these isolates to the species level is challenging. Recently, matrix-assisted laser desorption ionization time-of-flight mass spectrometry (MALDI-TOF MS) has been widely used for identification of most bacterial and fungal isolates. In this multicenter study, we evaluated the identification of *Nocardia* isolates using Autof MS1000 and Bruker Biotyper. A total of 86 non-duplicate *Nocardia* isolates from 7 hospital laboratories were evaluated. Further, we carried out sequence analysis of 16S rRNA, *gyrB, secA1, hsp65*, and *rpoB* genes as a reference method for *Nocardia* species identification. The 86 isolates were directly spotted on the target plate and plate protein extraction was performed. Data were analyzed by SPSS 19.0. In total, 72 (83.7%) strains (score ≥ 9.0) and 70 (81.4%) strains (score ≥ 2.0) were correctly identified by the Autof MS1000 and Bruker Biotyper systems, respectively, at the species level. There was no significant difference (P > 0.05) between the two systems using the same protein extraction method. In conclusion, the Autof MS 1000 and Bruker MALDI-TOF systems showed no difference in identification of *Nocardia spp*. to the species level and could meet the most important clinical requirement for species identification.

## INTRODUCTION

*Nocardia* species are slow-growing, gram-positive, branching, and weakly acid-fast bacilli belonging to the aerobic actinomycetes. *Nocardia* species are widespread in the environment and are recognized as the cause of serious human infections such as those affecting the lungs and brain (1, 2), especially in immunocompromised hosts. Currently, there are 137 recognized species in the *Nocardia* genus (see the List of Prokaryotic names with Standing in the Literature [http://www.bacterio.net/index.html]). These *Nocardia* species have different geographic prevalence, pathogenicity, and antimicrobial susceptibility patterns (3, 4). Therefore, it is important to identify *Nocardia* isolates to the species level for appropriate clinical treatment. For many years, phenotypic characteristics and biochemical methods were used for conventional identification. However, these methods are slow and insufficient to accurately distinguish clinical strains. With the development of molecular techniques, sequence analysis of several genes such as 16S rRNA, *gyrB, secA1, hsp65*, and *rpoB* can provide highly reliable results (5, 6). However, species-level identification is challenging. Currently, matrix-assisted laser desorption/ionization-time-of-flight (MALDI-TOF) mass spectrometry (MS) is a good choice for species-level identification of *Nocardia* (7–9). Two microbial MALDI-TOF MS identification systems, Bruker Biotyper MS (Bruker Daltonics, Bremen, Germany) and Vitek MS (bioMérieux, Marcy I’Etoile, France), have been used to identify *Nocardia* in clinical microbiology laboratories worldwide (8, 9). Notably, there is limited research on the performance of a Chinese MALDI-TOF MS-based system in identifying *Nocardia*. In this study, we compared the performance of the Autof MS 1000 system (Autobio Diagnostics, Zhengzhou, China) with that of the Bruker Biotyper MS system for identifying *Nocardia* species collected from seven clinical microbiology laboratories.

## MATERIALS AND METHODS

### Bacterial isolates

A total of 86 non-duplicate *Nocardia* isolates from seven Chinese clinical laboratories were evaluated. Among these *Nocardia* isolates, 43 were isolated from patients at Henan Provincial People’s Hospital from 2016 to 2020, 33 were isolated from Henan Chest Hospital from 2018 to 2020, 6 were isolated from Sixth People’s Hospital in Zhengzhou in 2019, and 4 strains were obtained from four other hospitals, including the Children’s Hospital Affiliated to Zhengzhou University, Central Hospital of Yellow River, Fuwai Cardiovascular Hospital of central China, and Xiping County People’s Hospital. All isolates were stored at −80°C before being transferred for two generations. After incubation at 35°C in air on Columbia blood agar with 5% sheep blood (Zhengzhou Antu Biological Co., Ltd., China) for 48 h to 72 h, these strains were used for MALDI-TOF MS identification.

### Reference method

We carried out the 5-locus (16S rRNA, *gyrB, secA1, hsp65*, and *rpoB*) multilocus sequence analysis (MLSA) as the reference method for *Nocardia* species identification. DNA extraction and amplification of all isolates were performed as described previously (5, 6). The criterion for molecular identification of *Nocardia* isolates was recommended according to the CLSI document, MM18-ED2 (10). For the genus *Nocardia*, the 16S rRNA sequence similarity of ≥ 99.6% was the criterion to identify an isolate at the genus and species level, and 99.0% to 99.5% was the criterion for identification at the genus level. All PCR products were sequenced on the ABI3730XL platform (Applied Biosystems, Foster City, CA, USA). The sequence similarity was obtained with the BLAST tool at the website of the National Center for Biotechnology Information (http://www.ncbi.nlm.nih.gov).

### MALDI-TOF MS identification

The bacterial isolates were subjected to two different treatments prior to MALDI-TOF identification: direct formic acid extractions on target and ethanol-formic acid extractions in tube. The differences lied in the sample processing procedures in the two methods. The turnaround time was 15 min for the first method and 45 min for the second method.

In the first method, we made slight changes to the commercial product manual; 1–2 single colonies were picked up with a 1 μL sterile loop, spotted onto a MALDI target plate, and dried. Consequently, the spots were overlaid with 1 μL of 70% formic acid twice. After the plate was dried, spectra were acquired and compared with the Biotyper software version 3.0 (db 4613; Bruker Daltonics, Billerica, MA, USA) and Autof MS (Autobio Diagnostics, Zhengzhou, China) databases.

The second method was performed as recommended by the manufacturer. For the Autof MS 1000 system, 1–2 single colonies were resuspended in 300 μL deionized water and 900 μL absolute ethanol in a 1.5 mL centrifuge tube, shaken, and mixed thoroughly. The suspension was centrifuged at 13,000 rpm for 3 min, and the supernatant was completely poured off. The precipitate was dried at 40°C for 3 min. A volume of 10 μL of 70% formic acid in water (Zhengzhou Antu Biological Co., Ltd., China) was added and mixed well, and 10 μL of 100% acetonitrile (Zhengzhou Antu Biological Co., Ltd., China) was subsequently added and mixed well. The suspension was centrifuged for 3 min at 13,000 rpm. One microliter of the supernatant was pipetted onto the MALDI target plate. For the Bruker Biotyper system, the tube-base extraction method was performed as recommended by the manufacturer. Briefly, the bacteria within the 1-μL sterile loop were resuspended in 300 μL of sterile water and 900 μL absolute ethanol. The suspension was centrifuged at 13,000 rpm for 2 min. The supernatant was discarded, and the pellet was resuspended in 25–100 μL of 70% formic acid in water and the same volume of 100% acetonitrile. After 2-min centrifugation at 13,000 rpm, 1 μL of the supernatant was spotted onto a polished steel MALDI target plate and then the protocol was continued as explained above.

The MALDI BIOTYPER 3.1 and Autof MS 1000 databases contain 35 (105 reference spectra) and 30 *Nocardia* species, respectively. The cut-off values for identification were established according to the manufacturer’s instructions. For the Autof MS 1000 system, score values >6.0 and <9.0 were considered identification at the genus level and ≥ 9.0 indicated identification at the species level. Score values ≤6.0 were considered unreliable identifications. For the Bruker Biotyper system, a spectral score of ≤ 1.70 was considered not to provide reliable identification, a score of >1.70 but < 2.00 indicated identification at the genus level, and a score of ≥2.00 was considered to indicate identification at the species level.

### Statistical analysis

The chi-square test was used to evaluate the performance of Autof MALDI-TOF MS and Bruker Biotype MS for the identification of *Nocardia* species. Data were analyzed using SPSS software package 20.0 (IBM; Chicago, IL, USA). A *P*-value below 0.05 was considered statistically significant for all analyses. Figures were generated using GraphPad Prism version 8.0 (GraphPad Software Inc., La Jolla, CA, USA).

### Reproducibility testing

Reproducibility testing was performed by two operators at each of three sites. A panel of five organisms (*N. cyriacigeorgica, N. farcinica, N. asteroides, N. wallacei*, and *N. otitidiscaviarum*) was tested in duplicate on two runs daily for five days. The identity of each organism was blinded with respect to the operators. Testing was performed using three different lot numbers for reagents. The position of each organism on the target slide was predetermined, and the organisms were tested sequentially on one slide and in a randomized manner on a second slide. Sample preparation, organism identification on the Autof MS 1000 system and Bruker Biotyper system v3.0, and result analysis were performed as described above.

## RESULTS

### Comparison of identification results of *Nocardia* species by Autof MS 1000 and Bruker Biotyper

Eighty-six *Nocardia* isolates from seven hospitals were collected in this study. Using MLSA, 11 species were identified, including *N. abscessus* (n=8), *N. asiatica* (n=2), *N. asteroides* (n=3), *N. beijingensis* (n=4), *N. brasiliensis* (n=3), *N. concave* (n=2), *N. cyriacigeorgica* (n=30), *N. farcinica* (n=22), *N. otitidiscaviarum* (n=8), *N. pseudobrasiliensis* (n=1), and *N. wallacei* (n=3). No significant difference in identification among species, genus, and nonidentification was observed between the two identification systems (Figure 1 and Table S1).

**Figure 1.**
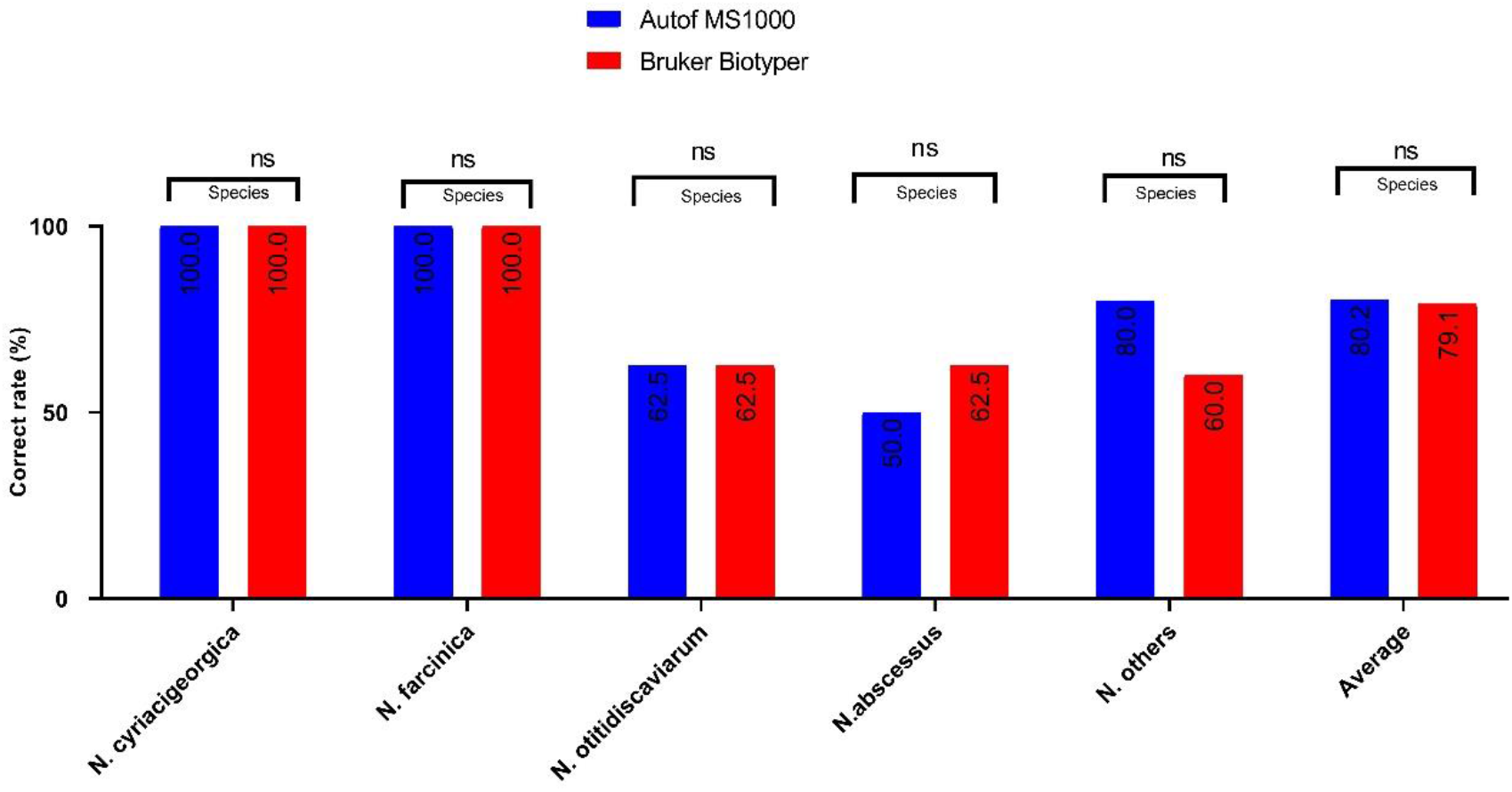
Comparison of identification results for *Nocardia* species between the Autof MS 1000 and Bruker Biotyper. ns: no significance.

With the direct formic extraction method, the identification accuracy rates of the Autof MS 1000 and Bruker Biotyper systems were 80.2% and 79.0% at the species level and 91.8% and 89.5% at the genus level, respectively. Using the ethanol-formic extraction method, the identification rates of Autof MS 1000 and Bruker Biotyper systems were 83.7% and 81.4% at the species level and 94.2% and 93.0% at the genus level, respectively. However, there were no significant differences found between the two systems at either the species or the genus level by the same protein extraction method (direct formic extraction method, P=0.923; ethanol-formic extraction method, P=0.913). No significant difference (Autof MS1000 system, P=0.838; Bruker Biotyper system, P=0.725) was found between the two different extraction processes in the Autof MS1000 or Bruker Biotyper systems (Figure 2 and Table S2).

**Figure 2.**
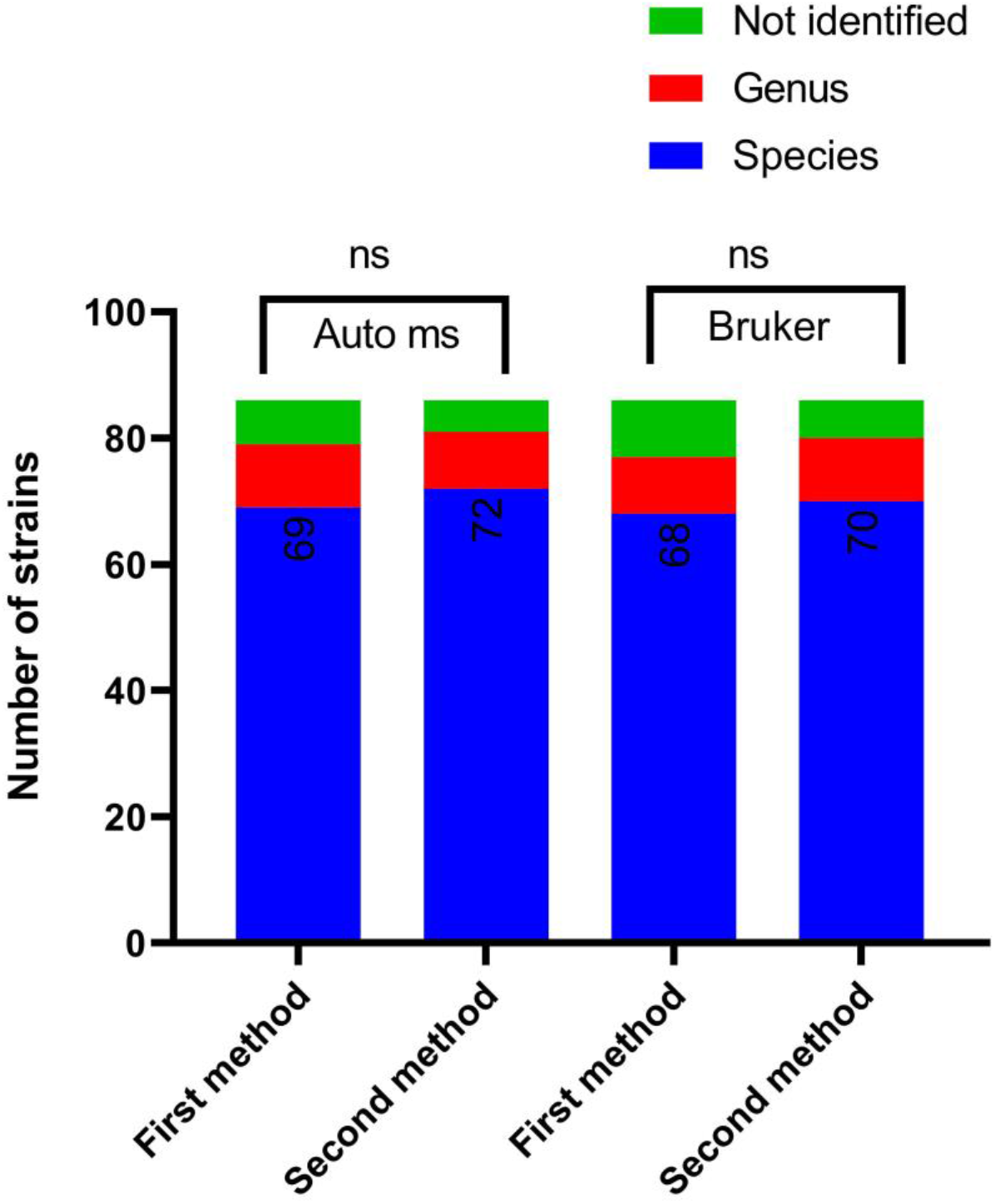
Comparison of the identification results of *Nocardia* species (n=86) by two different methods for the protein extraction protocol between the Autof MS and the Bruker Biotyper. ns: no significance.

### Incorrect identification results of *Nocardia* species by Autof MS1000 and Bruker Biotyper

Table 1 shows strains that could not be accurately identified. Two *N. abscessus* isolates could not be identified by both systems; two *N. otitidiscaviarum* isolates and one *N. wallacei* isolate could not be identified by the Autof MS1000 system, and two *N. wallacei* and two *N. beijingensis* isolates could not be identified by the Bruker Biotyper system.

**Table 1.**
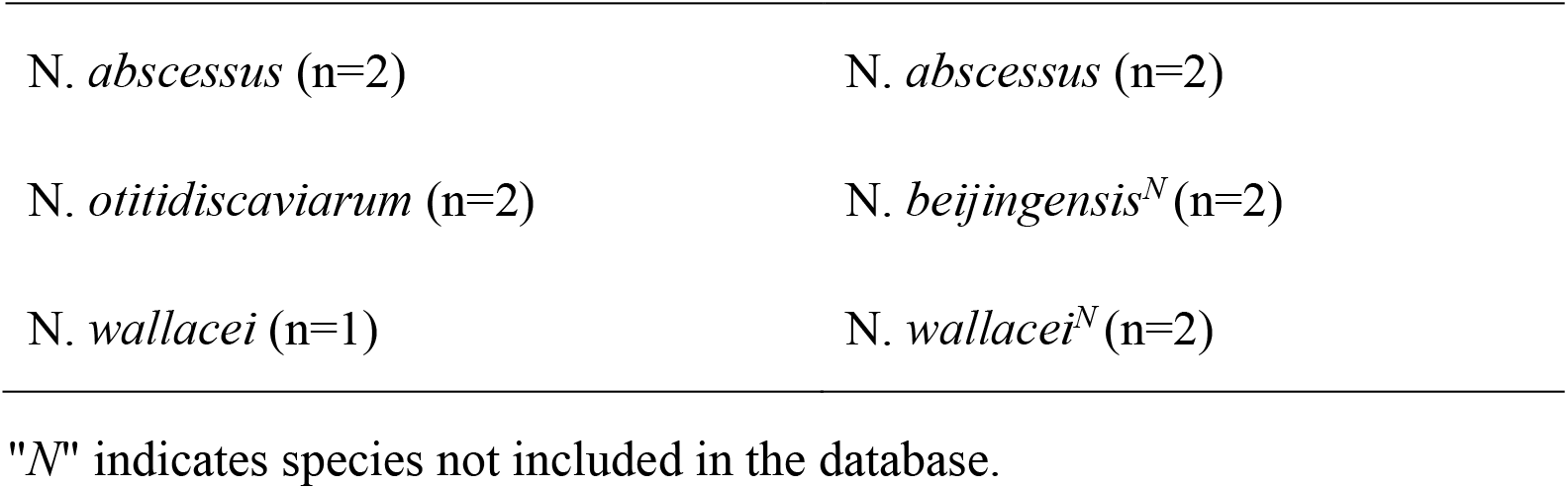
Incorrect identification of *Nocardia* by Autof MS 1000 and Bruker Biotyper MS.

## DISCUSSION

Due to the different drug-resistant phenotypes, the accurate identification of *Nocardia* species determines clinical precision treatment (11, 12). Molecular methodologies such as specific gene sequencing have emerged as the most accurate methods for identification of *Nocardia* (13, 14). However, sequencing is costly, has long turnaround times, and requires experienced personnel to circumvent technical traps; thus, it has only been implemented in reference laboratories or specialized centers (11). MALDI-TOF MS is a technology used for the identification of microorganisms (15, 16). It is a simple, reliable, and cost-effective method with a high degree of accuracy (17). This is especially important for routine clinical microbiology, as the results can be obtained directly from positive blood culture (18, 19), patient urine or other biological fluids (20, 21), as well as subcultures on agar plates and broth media (22, 23). Thus far, two MS identification systems, the Bruker Biotyper and Vitek systems, have been evaluated for identification of *Nocardia* strains (9, 24, 25). Here, we aimed to compare the commercial MALDI-TOF MS system for *Nocardia* identification (Bruker Biotyper) with two different methods of the Autof MS 1000 system from China. To the best of our knowledge, this is the first evaluation of the identification of *Nocardia* species by two Autof MS 1000 systems.

Generally, the MS system recommends two protein extraction methods commonly used in clinical laboratories, direct formic acid extractions on target and ethanol-formic acid extraction in tube, the first and the second method, respectively. The first method is simple, less time-consuming, and highly accurate for most bacteria. The second method is the standard operation method recommended by mass spectrometers, but it is difficult for laboratory personnel because it is a complex time-consuming process. The species-level identification accuracy for *Nocardia* using any extraction procedure was 36.4%–80.9% in Bruker Biotyper commercial databases (24, 25). We modified the protein extraction method to improve the accuracy of identification. During the *Nocardia*-wall lysing process, we added 1 μL of 70% formic acid cycle in the direct formic extraction method and extended the grinding time from 0.5 to 1 min in the ethanol-formic extraction method. We found that the modified protein-extraction method increased the species-level identification accuracy rates of *Nocardia* up to 79.0%–83.7% in the two systems.

In total, eight strains of *Nocardia* were misidentified, including two *N. abscessus*, two *N. beijingensis*, two *N. wallacei*, and two *N. otitidiscaviarum* strains. Due to slow growth, we could not collect enough protein profiles from the two *N. abscessus* strains. Since *N. beijingensis* was not included in the Bruker Biotyper database, but was included in the Autof MS 1000 database, the identification accuracy of the Autof MS 1000 system (100%, 4/4) was higher than that of the Bruker Biotyper system (50%, 2/4). Additionally, the Autof MS 1000 system could identify two *N. wallacei* to the genus level, benefiting from manual selection of the laser strike position to collect protein profiles due to the lack of *N. wallacei* and *N. beijingensis* in the databases of the Bruker Biotyper system. Moreover, two *N. otitidiscaviarum* strains were misidentified by the Autof MS 1000 system but were correctly identified by the Bruker Biotyper system. However, because of the limited number of strains, we could not demonstrate which system was more advantageous in the identification of *N. wallacei* and *N. otitidiscaviarum*, and more data are needed. Sample size (11 species) and study settings (the strains were from Henan Province located in the central area of China) were limiting factors in this study, as the research results only represented the experimental status of the local clinical epidemic isolates.

In summary, our multicenter study demonstrated that both the Autof MS 1000 and Bruker Biotyper systems were useful for the identification of clinical epidemic *Nocardia* species. The two systems showed no significant differences in species-level identification of *Nocardia*. We suggest that laboratories adopt modified protein-extraction methods to obtain better identification results of *Nocardia*.

## ACKNOWLEDGEMENTS

We thank Yunqi Tian (Xiping County People’s Hospital) for technical assistance and Yungang Han (Henan Chest Hospital), Lifang Ban (Sixth People’s Hospital of Zhengzhou city), Junwen Yang (Children’s Hospital Affiliated to Zhengzhou University), Xinkun Qi (Fuwai Cardiovascular Hospital of central China), and Changcheng Guo (Central Hospital of Yellow River) for providing additional clinical isolates. We thank the Department of Clinical Laboratory staff from Henan Provincial People’s Hospital (Bing Ma, Qiong Ma, and Shanmei Wang for conducting the experiments, and Youhua Yuan and Yi Li for helping to review the manuscript).

---

**Table S1.** Identification results of *Nocardia* by Autof MS 1000 and Bruker Biotyper.

| Identification by<br>MLSA (n=86) | Direct formic extraction method |  |  |  | Ethanol-formic extraction method |  |  |  |
| --- | --- | --- | --- | --- | --- | --- | --- | --- |
|  | Autof MS |  | Bruker |  | Autof MS |  | Bruker |  |
|  | 1000 |  | Biotyper |  | 1000 |  | Biotyper |  |
|  | specie | Genu | specie | Genu | specie | Genu | specie | Genu |
|  | s level | s | s level | s | s level | s | s level | s |
|  | level |  | level |  | level |  | level |  |
| *N. abscessus* (n=8) | 4 | 2 | 5 | – | 5 | 1 | 5 | 1 |
| *N. asiatica* (n=2) | – | 2 | – | 2 | – | 2 | – | 2 |
| *N. asteroides* (n=3) | 2 | 1 | 3 | – | 3 | – | 3 | – |
| *N. beijingensis*<br>(n=4) | 3 | 1 | – | 2 | 4 | – | – | 2 |
| *N. brasiliensis*<br>(n=3) | 3 | – | 3 | – | 3 | – | 3 | – |
| *N. concava* (n=2) | – | 2 | – | 2 | – | 2 | – | 2 |
| *N. cyriacigeorgica*<br>(n=30) | 30 | – | 30 | – | 30 | – | 30 | – |
| *N. farcinica* | 22 | – | 22 | – | 22 | – | 22 | – |
| (n=22) |  |  |  |  |  |  |  |  |
| *N. otitidiscaviarum* | 5 | – | 5 | 2 | 5 | 1 | 7 | 1 |
| (n=8) |  |  |  |  |  |  |  |  |
| *N. pseudobrasiliensis* | – | 1 | – | 1 | – | 1 | – | 1 |
| (n=1) |  |  |  |  |  |  |  |  |
| *N. wallacei* (n=3) | – | 1 | – | – | – | 2 | – | 1 |
| Total number of strains | 69 | 10 | 68 | 9 | 72 | 9 | 70 | 10 |

**Table S2.** Comparison of identification results of *Nocardia* with the Autof MS 1000 and Bruker Biotyper.

|  |  |  |  |  |  |  |  |
| --- | --- | --- | --- | --- | --- | --- | --- |
| Protein extraction protocol | Identification with Autof MS1000 |  |  | Identification with Bruker Biotyper |  |  | P value |
|  | species level | Genus level | Not identified | species level | Genus level | Not identified |  |
| Direct formic extraction | 69 | 10 | 7 | 68 | 9 | 9 | 0.923 |
| method |  |  |  |  |  |  |  |
| Ethanol-formic<br>extraction<br>method | 72 | 9 | 5 | 70 | 10 | 6 | 0.913 |
| P value | 0.838 |  |  | 0.725 |  |  |  |

